# Predicting range shifts of three endangered endemic plants of the Khorassan-Kopet Dagh floristic province under global change

**DOI:** 10.1101/2020.09.19.304766

**Authors:** Mohammad Bagher Erfanian, Mostafa Sagharyan, Farshid Memariani, Hamid Ejtehadi

**Affiliations:** Quantitative Plant Ecology and Biodiversity Research Lab., Department of Biology, Faculty of Science, Ferdowsi University of Mashhad, Mashhad, Iran; Department of Plant Biology, Faculty of Biological Science, Tarbiat Modares University, Tehran, Iran; Department of Botany, Research Center for Plant Sciences, Ferdowsi University of Mashhad, Mashhad, Iran

**Author notes:** Corresponding author: M.B. Erfanian, PO BOX 9177948974.

**Keywords:** Climate change, Khorassan-Kopet Dagh, Binalood Mountains, Endemic plants

## Abstract

Khorassan-Kopet Dagh (KK) floristic province is an ecoregion that has rarely been studied. A total number of 2576 (356 endemic) vascular plants have been recorded from this area. Most of the endemic species of KK are rare and range-restricted. To assess the vulnerability of plant species under a rapidly changing climate, we can use species distribution modelling (SDM) to predict their potential present and future distribution. We used SDM to evaluate range size changes for three (critically) endangered endemic species to KK, namely *Nepeta binaloudensis, Phlomoides binaludensis*, and *Euphorbia ferdowsiana*. These plants represent KK endemic species that grow in the different elevation ranges of KK mountains. Using the HadGEM2-ES general circulation model and two Representative Concentration Pathways Scenarios (RCP), including RCP 2.6 (most optimistic) and RCP 8.5 (most pessimistic), we predicted the potential present and future (i.e., 2050 and 2070) distribution for each species. The ensemble model of nine different methods was used for this prediction. Our results showed that, except for *P. binaludensis* that will face range contraction, the other species would benefit from potential range expansion in the future. *Euphorbia ferdowsiana* will remain limited to a narrow range of KK. However, the other two plants will have suitable habitats in various mountains of KK. To conserve flora of KK, we urge using management efforts with a focus on providing elevational migration routes at the local scales in KK mountains. Additionally, assisted migration among different mountains of this region will be beneficial to conserve its endemic plants. For *E. ferdowsiana* genetic diversity storage employing seed banks and botanical garden preservation should be considered.

## 1 Introduction

Geographical distributions of plant species are mainly delimited by climatic conditions ^1^. There is a considerable amount of research declaring climate change lead to pole or upward changes in this species distribution in the past decades (e.g. ^2, 3^, and ^4^). To assess the vulnerability of plant species under a rapidly changing climate, we can use species distribution modelling (SDM) to predict the climate niches of them and project their potential future shifts in their spatial distributions ^5^. The SDM approach results could be used to develop adaptive management strategies, including assisted migration to ensure that species will compensate climate change effects ^1,2,6^.

Khorassan-Kopet Dagh (KK) floristic province is an under-studied ecoregion that is located mainly in northeastern Iran, and partly in southern Turkmenistan and northwestern Afghanistan (Figure 1) ^7,8^. A total number of 2576 of vascular plant species have been recorded from this region. Among these, 356 species are endemic to KK ^9^. *Nepeta binaloudensis* Jamzad (Lamiaceae) is a perennial species endemic to KK. This plant grows in the 2300 to 3000 m a.s.l. elevational range in the Binalood Mountains and it has recently been recorded from the Hezar-Masjed Mountains ^10–12^. This species was assessed as an Endangered plant in KK ^13^. *Phlomoides binaludensis* Salmaki & Joharchi (Lamiaceae) is another endemic species to KK. Hitherto, *P. binaludensis* is recorded only from the Binalood Mountains of KK. This perennial plant grows in an elevation range between 1350 and 2000 m a.s.l. This species was categorised as an Endangered plant ^13^. *Euphorbia ferdowsiana* Pahlevani is an endemic perennial plant growing in the eastern slopes of the Binalood Mountains. It has a narrow distribution range in the Binalood Mountains and has been recorded from an elevation range of 2100 to 2700 m a.s.l. ^14^. Because of its limited distribution range, this plant was categorised as a Critically Endangered species ^13^.

**Figure 1.**
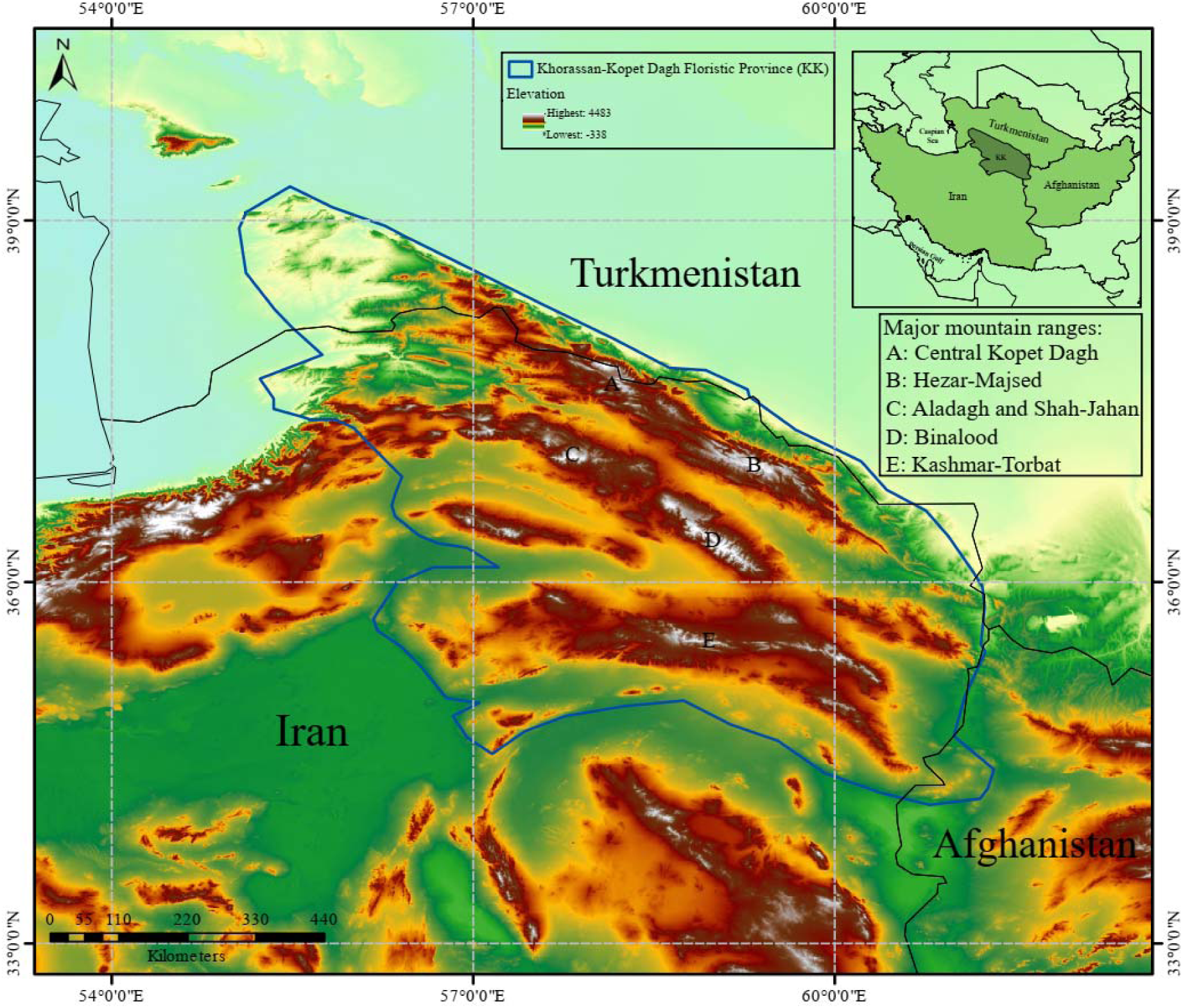
Study area – Khorassan-Kopet Dagh floristic province (KK). The reported elevation range in the legend is for the entire map and not corresponds to the elevation range of KK. The original figure was adopted from Memariani (2020).

Here, considering three (critically) endangered endemic species from KK that grow in an elevation range of 1350 to 3000 m a.s.l., we aim to evaluate how climate change affect their distribution. Specifically, our goals are to (a) predict the potential present distribution of these threatened species; (b) study the effects of climate change on the distribution of these plants by using an ensemble model from nine different modelling approach for 2050 and 2070 under the most optimistic and pessimistic scenarios; (c) evaluate the climate change-induced elevational range shifts of these plants.

## 2 Materials and methods

### 2.1 Study area

Our study area was Khorassan-Kopet Dagh (KK) ecoregion (Figure 1). This mountainous region, with more than 165,000 km^2^ area, has a complex topography ranging from *ca*. 250 m in lower foothills to the elevations higher than 3000 m in the Binalood, Hezar-Masjed, and Shah-Jahan Mountains ^8,9^. KK has a continental climate. The mountainous ranges of KK have a Mediterranean or Irano-Turanian xeric-continental bioclimate with an average annual precipitation of 300-380 mm. The mean annual temperature in the area shows elevation-dependent values and ranges from 12°C to 19°C ^8,15^. KK is home to diverse vegetation types, and among them, the montane steppes and grasslands are the most abundant. The area is hosting 2576 species or infraspecific taxa. Among these species, 356 (13.8 per cent of the total species pool) are endemic to this ecoregion ^8,9,13^. Most of the endemic species of KK are range-restricted and rare ^16^. The mountainous habitats of KK are threatened by various disturbances (e.g. overgrazing, land-use change, recreation activities). Only *ca*. 8 per cent of KK habitats are protected and managed with various protection guidelines ^8,9,17–21^.

### 2.2 Species data

Occurrence records for all three species were obtained using direct observation from field surveys, including the records from Ferdowsi University of Mashhad Herbarium (FUMH). These surveys were mostly one-time field surveys to collect plant specimens from different parts of KK. Most of the data collection was carried out from 2002 to 2018. These occurrence points were recorded from the whole distribution range of the studied species. Absence points were collected from plot data or field surveys. Due to its collection of from the Hezar-Masjed Mountains, we opted to use only plot data as the absence points of *N. binaloudensis* to avoid adding uncertain data for the modelling. However, for the other species, as they were endemic to the Binalood mountains, we included the absence points from field surveys. Additionally, for these two species, only plot data were considered for recording true absences from the Binalood Mountains.

The species occurrence data were spatially thinned by randomly selecting a present point within a single grid cell of the predictor variable layer using the spThin package ^22^. Three sets of background points (n=10000) were generated using the biomod2 package ^23^ to be used in the SDM. Finally, for *N. binaloudensis*, 13 presence points + 36 true absence points + 3 * 10000 background points; for *P. binaludensis*, 11 presence points + 569 true absence points + 3 * 10000 background points; and for *E. ferdowsiana*, 5 presence points + 580 true absence points + 3 * 10000 background points were used for the modelling.

### 2.3 Environmental data

We used physiographic maps and bioclimatic variables as the environmental data in SDM. Physiographic maps were: elevation, slope, and aspect. In this study, 19 bioclimatic layers (table 1) which are reliable in defining the physio-ecological tolerances of a species were used. These layers downloaded from Worldclim ^24^. In addition to present layers (i.e., those of the year 2000), future layers for 2050 and 2070 were also download. For downloading future layers, the Hadley Centre Global Environmental Model version 2▫Earth System (HadGEM2-ES) general circulation model (GCM) and two Representative Concentration Pathways (RCPs) scenarios, including RCP 2.6 (most optimistic) and RCP 8.5 (most pessimistic) were selected. The species distribution models obtained by using these data were evaluated to assess climate change impacts on the distribution and elevational shifts of the studied species.

**Table 1.** The nineteen bioclimatic variables that were used for modelling in the present study.

| Bioclimatic variable | name |
| --- | --- |
| Mean annual temperature | BIO1 |
| Mean Diurnal Range | BIO2 |
| Isothermality | BIO3 |
| Temperature Seasonality | BIO4 |
| Maximum Temperature of Warmest Month | BIO5 |
| Minimum Temperature of Coldest Month | BIO6 |
| The annual temperature range (BIO5-BIO6) | BIO7 |
| the mean temperature of wettest quarter | BIO8 |
| the mean temperature of the driest quarter | BIO9 |
| Mean temperature of warmest quarter | BIO10 |
| Mean temperature of coldest quarter | BIO11 |
| Annual Precipitation | BIO12 |
| Precipitation of Wettest Month | BIO13 |
| Precipitation of Driest Month | BIO14 |
| Precipitation Seasonality (standard deviation/mean) | BIO15 |
| Precipitation of Wettest Quarter | BIO16 |
| Precipitation of Driest Quarter | BIO17 |
| Precipitation of Warmest Quarter | BIO18 |
| Precipitation of Coldest Quarter | BIO19 |

To remove collinear layers (i.e., physiographic and bioclimatic layers), we perform a Principal Component Analysis (PCA). The PCAs were performed in the ade4 package ^25^. Then, layers with the most contribution in explanation of the variation in the species presence points space were selected. This analysis was performed as suggested by Guisan, Thuiller, and Zimmermann (2017). Moreover, the variance inflation factor (VIF) for the selected layers was calculated by performing a step-by-step process using the usdm package ^27^. Finally, we selected those variables with VIF <10. A list of the selected variables for each species is presented in table S1 of supplementary data.

### 2.4 Modelling settings

Nine modelling methods, including the Generalised Linear Model (GLM), Generalized Additive Model (GAM), Generalized Boosting Model (GBM), Classification Tree Analysis (CTA), Artificial Neural Network (ANN), Surface Range Envelop (SRE, also known as BIOCLIM), Multiple Adaptive Regression Splines (MARS), Random Forest (RF), and Maximum Entropy (MAXENT), were used in this study. These nine methods are available in the biomod2 package. We randomly split the occurrence data into two subsets, 70 per cent of the data was used for the model calibration, and the remaining 30 per cent was used for the model evaluation. We split data because we had no independent data for the model evaluation. The number of replications for each model calculation was set to ten.

To measure SDM performances, we employed the True Skill Statistic (TSS) and Area Under Curve (AUC) of the Receiver Operation Curves (ROCs). TSS is a threshold-dependent measure, ranges from -1 and +1, where +1 indicates perfect agreement between predictions and observations, and values of 0 or less indicate agreement no better than random partitioning ^28^.

AUC is widely used to determine the predictive accuracy of SDMs. Generally, AUC ranges from 0.5 to 1.0 and models with AUC > 0.9 are categorised as very good ^4^. For the binary transformation, we employed the threshold maximises TSS to convert the occurrence probability values into presence/absence predictions. The thresholding approach maximising the TSS is well suited since it produces the same threshold using either presence-absence data or presence-only data ^26^. These calculations were performed using the biomod2 package ^23^.

### 2.5 Ensemble forecasting

We used the ensemble forecasting procedure to obtain final models in order to reduce the uncertainty among the species distribution methods. To combine models, those with TSS > 0.9 were selected. Ensemble models were predicted for current and future conditions at a 1-km^2^ resolution. Then, the ensemble models were converted into binary presence-absence predictions using the threshold that maximises TSS. The ensemble forecasting was performed in the biomod2 package.

### 2.6 Range and elevational shifts

To evaluate the range size changes of all three species, we compared the predicted models for the future distribution of the species to that of the present. Then, we distinguished four distinct habitat types; (a) stable habitats: habitats in which species is present and predicted to be present in the future; (b) lost habitats: habitats in which species is present, but climate change will lead to local extinction of the species in them; (c) gained habitats: habitats in which species is not present, but climate changes will make them suitable for species growth; (d) unsuitable habitats: habitats that are unsuitable for species both in the present and future climatic conditions. Range size changes were predicted using the biomod2 package ^23^.

To compare elevational shifts for each species, we extracted the elevation of the grids correspond to species present from the current and future ensemble models. Then, density plots comparing the elevational range of the present and future habitats were drawn in R. All calculations were performed in R ver. 3.6 ^29^.

## 3 Results

### 3.1 Modelling evaluation

The MAXENT and GLM methods showed a better performance than the other methods. The ensemble models had the best overall performance with a TSS and ROC>0.99. The predictive performances of the nine modelling methods for all three species showing inter-model variability are presented in Figure 2.

**Figure 2.**
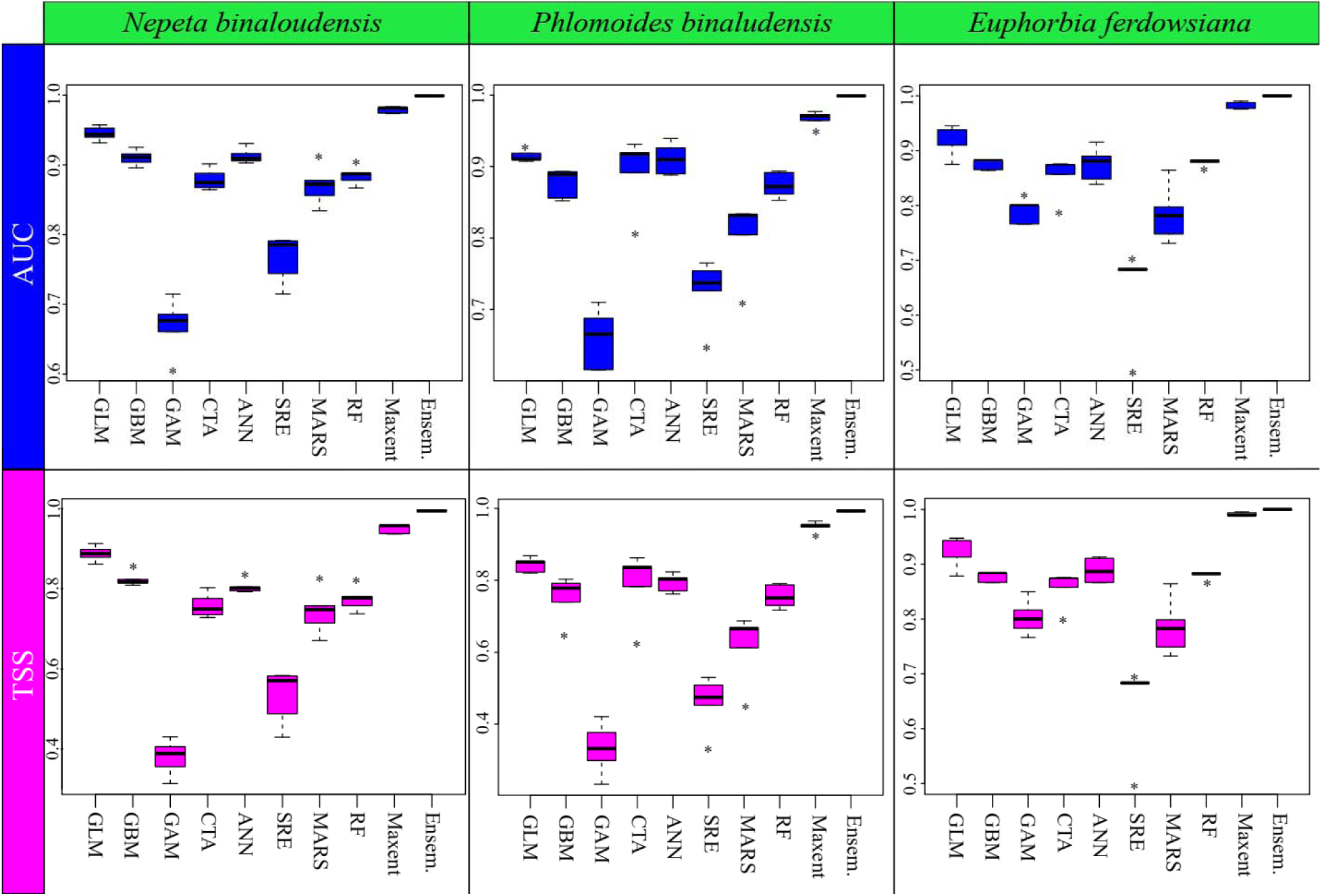
Boxplots showing the AUC and TSS scores for model evaluation from ten cross▫validation runs on test data for the nine SDM methods that were used for the prediction of three species distribution in the Khorassan-Kopet Dagh floristic province. For comparison, the evaluation scores for the ensemble model are shown. The ensemble model does not include those models with a TSS < 0.9. See the text for model abbreviations

### 3.2 Impacts of climate change on *Nepeta binaloudensis*

The ensemble habitat suitability map showed that the area of the currently suitable habitats for *N. binaloudensis* was 3407 km^2^ (Figure 3). Note that the base map that was used for showing SDM results does not match that of KK boundaries and also covers peripheral areas of KK. The range size analysis showed that in 2050 and under RCP 2.6 (the most optimistic) scenario, 708 km^2^ of the currently suitable habitats will be lost, which is approximately 21 per cent of the current habitats. However, 3408 km^2^ will become newly suitable habitats for this species that is approximately 100 per cent increase of the current range size. Finally, 2699 km^2^ of the current habitats will remain suitable for this species. Thus, the range size change for this species will be 79 per cent. In 2050 and under RCP 8.5 (the most pessimistic) scenario, 963 km^2^ (28 per cent) of the currently suitable habitats will become unsuitable; on the other hand, 2947 km^2^ will become suitable for the species distribution – 86 per cent increase of the current range size. Finally, 2444 km^2^ (72 per cent) of the current habitat size will remain suitable. As a result, the changes in the range size for this species will be 58 per cent (Figure 4).

**Figure 3.**
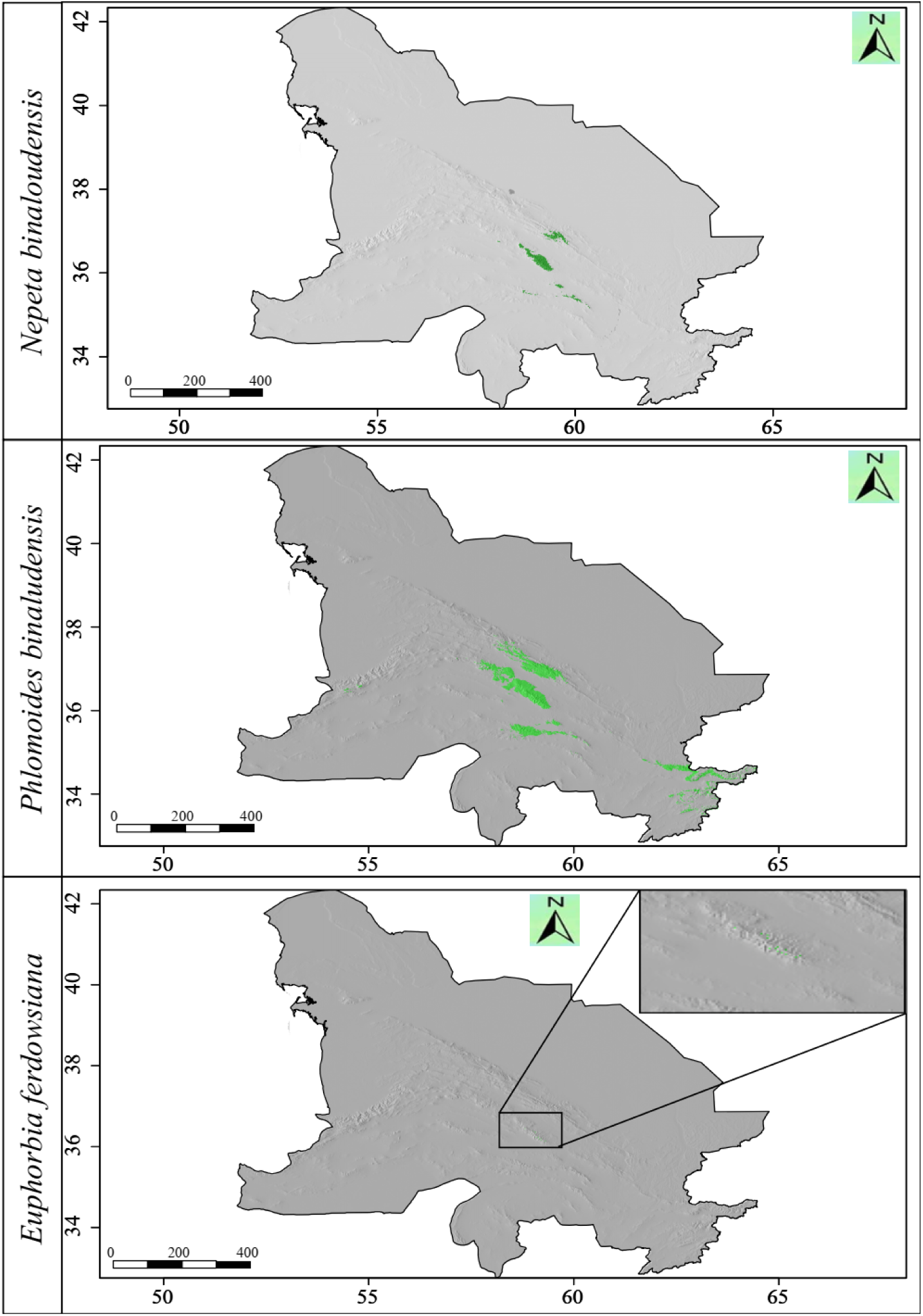
The habitat suitability maps of *Nepeta binaloudensis, Phlomoides binaludensis*, and *Euphorbia ferdowsiana* under the current climate conditions in the Khorassan-Kopet Dagh floristic province. Green areas indicate suitable habitats. The base map does not match that of KK boundaries and covers peripheral areas of KK.

**Figure 4.**
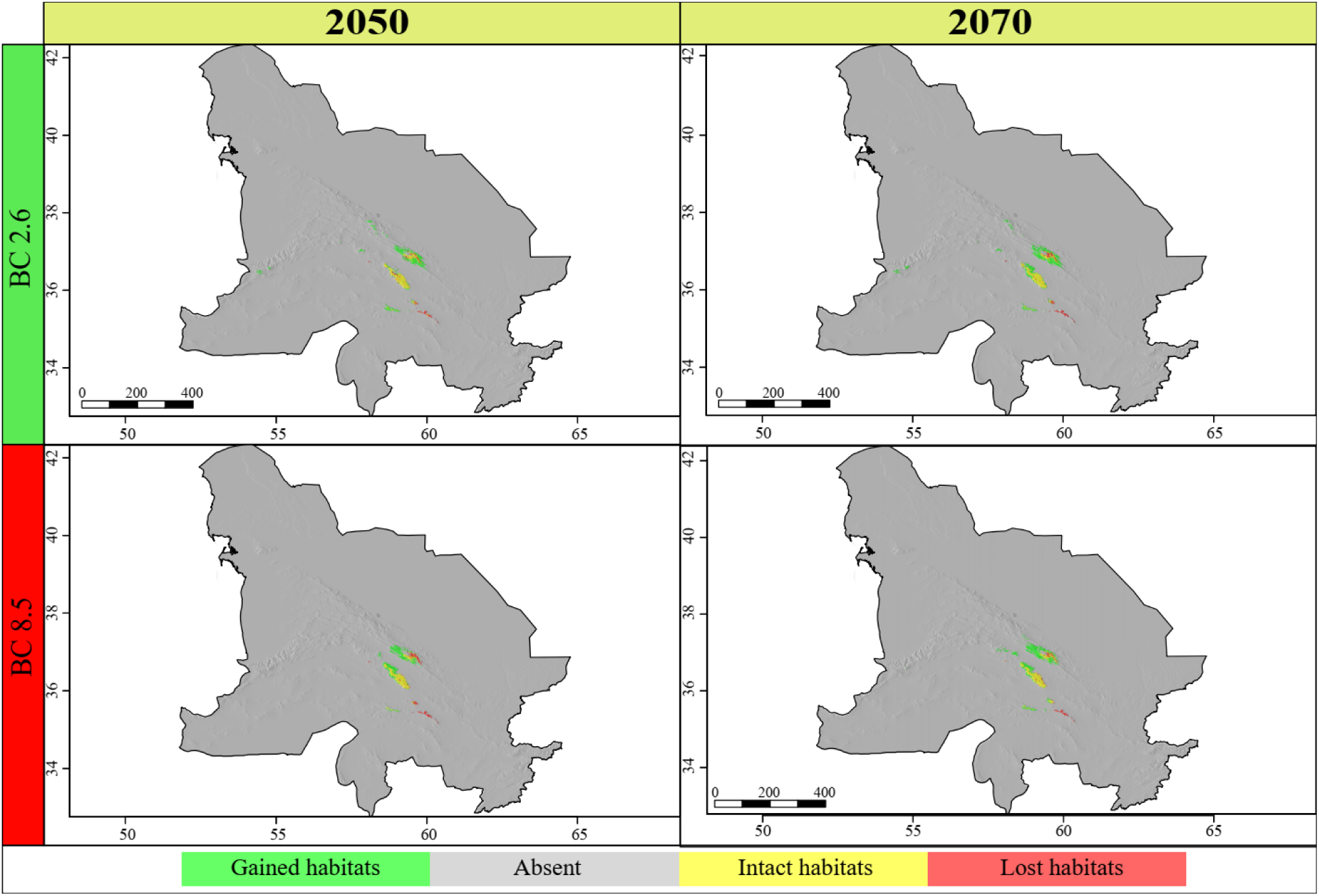
Range size changes of *Nepeta binaloudensis* in the future years (2050 and 2070) under the most optimistic (RCP 2.6) and pessimistic (RCP 8.5) scenarios. The base map does not match that of KK boundaries and covers peripheral areas of KK.

In the 2070 and under RCP 2.6 scenario, 771 km^2^ (23 per cent) of the currently suitable habitats will be lost, 2636 km^2^ (77 per cent) will remain suitable, and 4333 km^2^ will be newly-suitable areas for this species which is 127 per cent increase in distribution. The range size changes for this plant will be 105 per cent. Under RCP 8.5 scenario in 2070, 680 km^2^ (20 per cent) will be lost. The stable area covers 2727 km^2^ (80 per cent), and 4096 km^2^ will become suitable for this species growth (i.e., 120 per cent habitat gain) (Figure 4). Therefore, the range size change will be 100 per cent for this year and under this scenario.

The elevation shift for *N. binaloudensis* in 2070 and 2050 will be downslope as this species will grow in the lower elevations. Under RCP 2.6, a potential bimodal presence might be detected. The optimum elevational range for this species growth will be decreased under RCP 8.5 (Figure 5).

**Figure 5.**
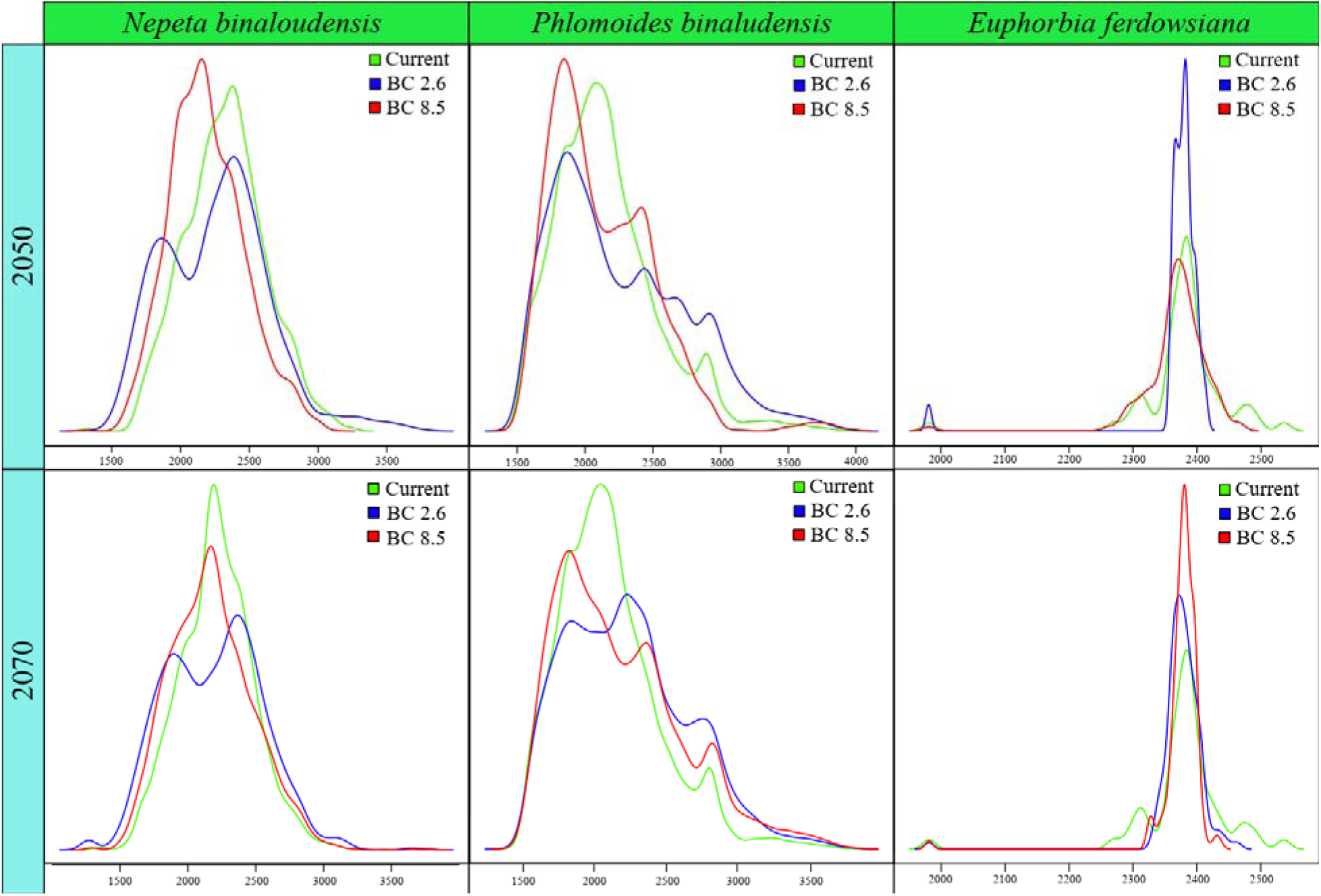
Density plots showing the elevation of the current and future suitable habitats of all three species.

### 3.3 Impacts of climate change on *Phlomoides binaludensis*

The ensemble habitat suitability map showed that the current range size (i.e., currently suitable habitats) for *P. binaludensis* was 19311 km^2^ (Figure 3). In 2050 and under RCP 2.6 scenario, 13733 km^2^ (71 per cent) of the currently suitable habitats will be lost, 5578 km^2^ (29 per cent) will remain suitable, and 1441 km^2^ will be newly-suitable areas for this species – 7 per cent increase in distribution. The range size changes for this plant will be -64 per cent. Under RCP 8.5 scenario in 2050, 14126 km^2^ (73 per cent) will be lost. The stable habitats cover 5185 km^2^ (27 per cent), and 203 km^2^ will become suitable for this species growth – 1 per cent habitat gain. As a result, the range size changes for this species will be -72 per cent (Figure 6).

**Figure 6.**
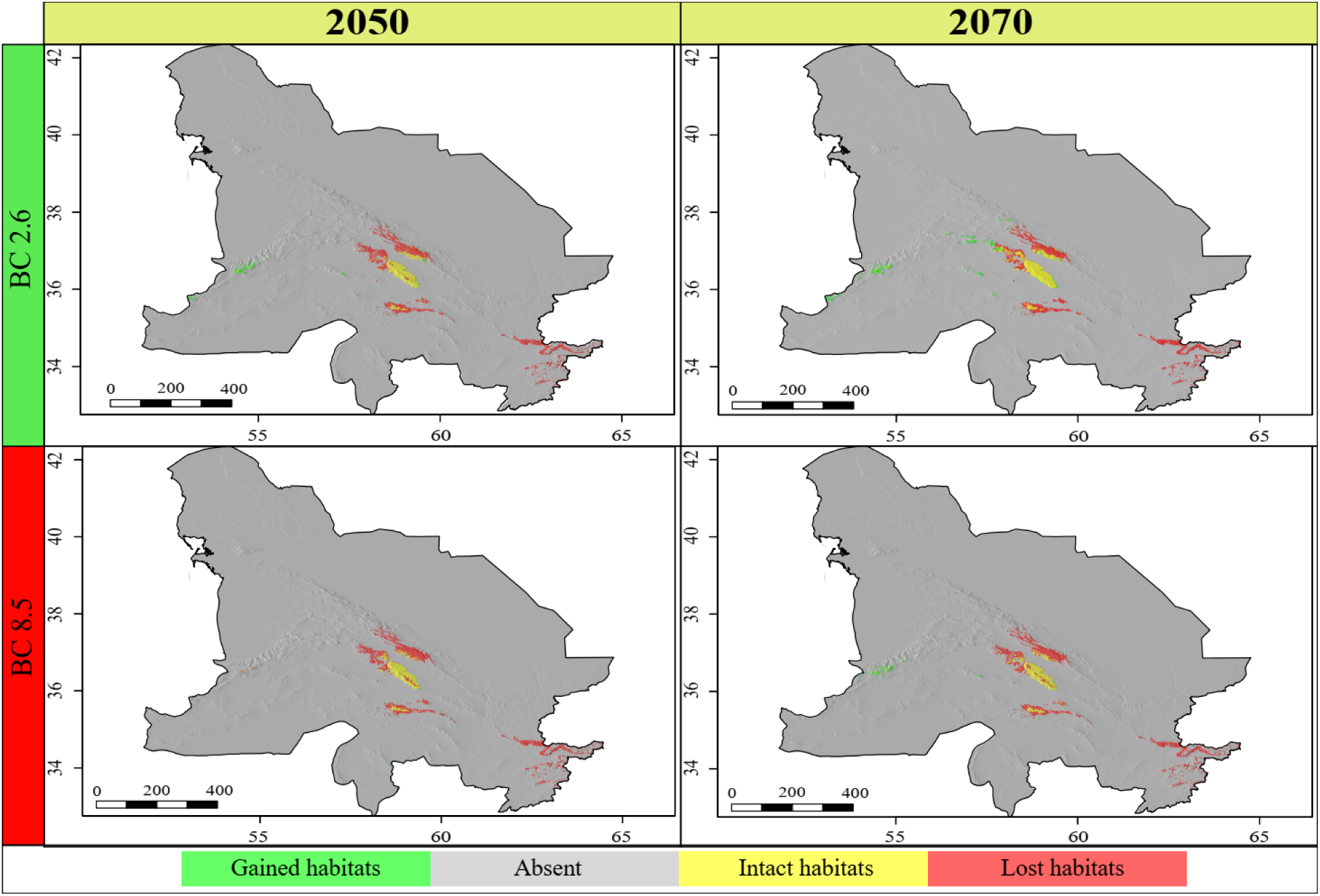
Range size changes of *Phlomoides binaludensis* in the future years (2050 and 2070) under most optimistic (RCP 2.6) and pessimistic (RCP 8.5) scenarios. The base map does not match that of KK boundaries and covers peripheral areas of KK.

In 2070, under RCP 2.6 scenario, 12573 km^2^ (65 per cent) of the current habitats will become unsuitable, on the other hand, 2877 km^2^ will become suitable for this species growth – 15 per cent habitat gain. Also, 6738 km^2^ (35 per cent) of the current habitat size will remain suitable. As a result, changes in the range size for this species will be -50 per cent. In this year and under RCP 8.5 scenario, 13892 km^2^ (72 per cent) of the currently suitable habitats will be lost, 5419 km^2^ (28 per cent) will remain suitable, and 1070 km^2^ (i.e., 6 per cent habitat gain) will be newly-suitable areas for this species. The range size changes for this species will be -66 per cent (Figure 6).

Climate change will induce an elevational shift in *P. binaludensis*. This shift will generally be a downslope. The current optimum elevational range for this species is 2000-2300 m a.s.l. Climate change mostly influences habitats in this optimum range, and this species will force to invade lower elevations (Figure 5).

### 3.4 Impacts of climate change on *Euphorbia ferdowsiana*

The ensemble habitat suitability map showed that the current range size (i.e., currently suitable habitats) of *E. ferdowsiana* was 60 km^2^ (Figure 3). In 2050 and under RCP 2.6 scenario, 47 km^2^ (78 per cent) of the currently suitable habitats will be lost, 13 km^2^ (22 per cent) will remain suitable, and 29 km^2^ (48 per cent) will be newly-suitable habitats for this species. Therefore, the range size changes for this species will be -30 per cent. Under RCP 8.5 scenario in 2050, 39 km^2^ (65 per cent) will be lost. The stable habitats cover 21 km^2^ (35 per cent), and 108 km^2^ will become suitable for this species growth – 180 per cent increase in distribution. As a result, the range size changes for this species will be 115 per cent (Figure 7).

**Figure 7.**
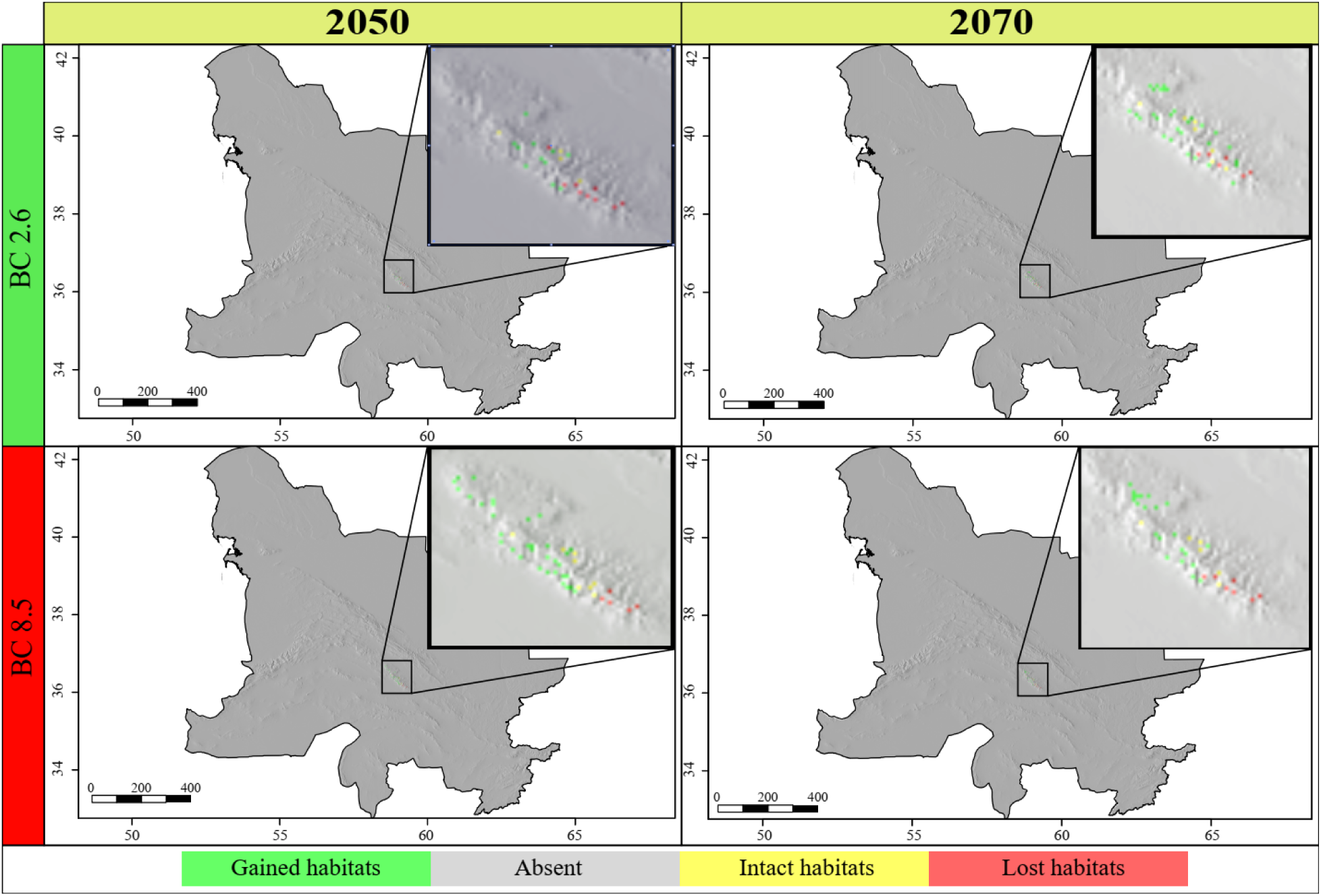
Range size changes of *Euphorbia ferdowsiana* in the future years (2050 and 2070) under most optimistic (RCP 2.6) and pessimistic (RCP 8.5) scenarios. The base map does not match that of KK boundaries and covers peripheral areas of KK.

In 2070, under RCP 2.6 scenario, 36 km^2^ (60 per cent) of the current habitats will become unsuitable, on the other hand, 93 km^2^ (155 per cent) will become suitable for this species growth. Also, 24 km^2^ (40 per cent) of the current habitat size will remain suitable. As a result, changes in the range size for this species will be 95 per cent. In this year and under RCP 8.5 scenario, 44 km^2^ (73 per cent) of the currently suitable habitats will be lost, 16 km^2^ (27 per cent) will remain suitable, and 75 km^2^ will be newly-suitable areas for this species (i.e., 125 per cent habitat gain). The range size changes for this species will be 52 per cent (Figure 7).

Climate change (under both scenarios in 2070 and 2050) will affect the distribution of this species in the elevation gradients. As a result, the suitable elevational range of this species will become narrow with an optimum of 2400 to 2600 m a.s.l (Figure 5).

## 4 Discussion

In the current study by evaluating three (critically) endangered species endemic to KK mountains, we showed that climate change will lead to species specific range expansion/contradiction and elevational shifts. Our results also revealed the variability among different SDM methods. Since there is no best model for species distribution modelling, we used an ensemble model of the nine different models. The ensemble model outperformed the other models. The efficiency of ensemble models was also reported by ^5,30,31^. However, Hao et al. (^32^) found no particular benefit in using ensembles over individual models.

### 4.1 How climate change shape the distribution of *Nepeta binaloudensis*?

The ensemble map showing the current distribution of *N. binaloudensis* indicated that, with two patches in the Hezar-Masjed and Kashmar-Torbat Mountains, this species is mainly stablished in the Binalood Mountains. Hitherto, this plant has not been recorded from the Kashmar-Torbat Mountains. The analysis on the effects of climate change on this species in 2050 and 2070 under the most optimistic scenario (RCP 2.6) showed that most parts of the currently suitable habitats that located in the Binalood Mountains will remain unaffected. While range expansion will be happening in the Hezar-Masjed Mountains in 2050 and continue in 2070, the currently suitable habitats in the Hezar-Masjed Mountains will become unsuitable. Note that this species has not been recorded from the predicted lost habitats in the Kashmar-Torbat Mountains. In 2050 and 2070, a northwestern migration trend will be detected. As a result, habitats in the Central Kopet Dagh and Aladagh and Shah-Jahan Mountains will become suitable for this plant. Under the most pessimistic scenario (RCP 8.5) in 2050 and 2070, a similar migration trend of RCP 2.6 will be detected. However, the Central Kopet Dagh Mountains will not become suitable habitat for this plant. In general, as a result of climate change, a range expansion trend will be observed for this species. Range expansion as the result of climate change impacts was reported for endemic herbs of the flora of Namibia ^33^. Macel et al. ^34^ suggested that range expanding plants evolve an increased vigour or an altered defence system for adaption to new habitats. Our results indicated, as a response to climate change impacts, this species will migrate to the lower elevations. Alavi et al. ^5^ reported that *Taxus baccata* will go upslope as a response to climate change.

### 4.2 How climate change shape the distribution of *Phlomoides binaludensis*?

The ensemble model for this species predicted that as well as the Binalood Mountains, *P. binaludensis* suitable habitats are found in the Hezar-Masjed, Aladagh and Shah-Jahan, and Kashmar-Torbat Mountains. Furthermore, some parts of the northwestern mountains of Afghanistan are the potential suitable habitats for this plant. So far, from the extensive field surveys in KK, this species has only been found in the Binalood Mountains. According to our results (Figure 3), in the Binalood Mountains, this species has an broad distribution range, suggesting the least vulnerability of its current habitats.

Due to the climate change in both scenarios and years, great range contraction will be observed for this species. Considering the limited range of this plant, naturally, it could not migrate to the predicated gained habitats across KK. Consequently, assisted migration should be used to introduce it to newly suitable habitats. In the Binalood Mountains, newly suitable habitats will be found in the southeastern slopes. In this mountain range, similar to *N. binaloudensis*, this species will experience an elevational shift towards lower elevations. Furthermore, populations of this plant which grow in higher elevations of the Binalood Mountains will become extinct. Range contraction has been reported for many species (e.g., ^35, 36, 37, 38^, and ^5^).

### 4.3 How climate change shape the distribution of *Euphorbia ferdowsiana*?

The ensemble map of the current distribution of *E. ferdowsiana* showed that the currently suitable habitats for this plant are restricted to the Binalood Mountains. The southern parts of this mountain range are the favourable habitats for this species. The limited current distribution range of the plant supports categorising it as a critically endangered species. The effects of climate change on this species in 2050 and 2070 under the most optimistic scenario (RCP 2.6) is that current habitats will become unfavourable and this species should migrate to the northern parts of the Binalood Mountains. Under the most pessimistic scenario (RCP 8.5), a northwestern migration trend also will be detected and the current southern populations will become extinct. Generally, this critically endangered species will experience a limited range expansion within the Binalood Mountains. This study results showed that this species has a narrow optimum elevational range. Due to climate change, this narrow range will become narrower. Moreover, habitats that are located in this elevational range in the Binalood Mountains faces intensive grazing by sheep and goats ^18^.

### 4.4 A note on the conservation of these species

Our results indicated that no significant extinction risk will threaten *N. binaloudensis*, as this species, on average, could benefit from 85 per cent increase in its distribution range size considering all the models and years. On the other hand, *P. binaludensis*, on average will lose 63 per cent of its current suitable. Hence, climate change will greatly affect this species. This finding should be considered in future conservation programs. These two plants will force to grow in lower elevations of the Binalood Mountains. However, habitats detected as their potential suitable ranges are under various disturbances (e.g. land use changes, overgrazing, recreation) ^17^. *Euphorbia ferdowsiana -* with a limited distribution in high elevations of the Binalood Mountains - will benefit from 58 per cent range expansion. *Euphorbia ferdowsiana* should be introduced to the northern parts of the Binalood Mountains to ensure its survival because this plant will lose most of its current habitats.

## 5. Conclusions

We studied three endangered plants that grow in different elevational ranges of KK mountains. Except for *P. binludensis* that will experience a southern migration trend, the other species will migrate in with a northward trend. This study results showed that beside range contraction endangered species also could gain favourable habitats in the future. Our results can be used to propose proper management and conservation of the endangered species in the under-studied KK region. Our results suggested that ensuring a successful migration for an endangered species could be a proper strategy for conserving threatened species from climate change impacts.

## Acknowledgements

The authors wish to thank the Ferdowsi University of Mashhad for the financial support of this research.

## Author contributions

MBE, HE, and MS designed the study. FM and MS provided the data. MBE analysed the data and prepared figures and tables. MBE and MS write the first draft. The final draft was approved by all of the authors.

## Data archiving

The data regarding this study will be published online.

